# Searching more genomic sequence with less memory for fast and accurate metagenomic profiling

**DOI:** 10.1101/036681

**Authors:** Shea N. Gardner, Sasha K. Ames, Maya B. Gokhale, Tom R. Slezak, Jonathan E. Allen

## Abstract

Software for rapid, accurate, and comprehensive microbial profiling of metagenomic sequence data on a desktop will play an important role in large scale clinical use of metagenomic data. Here we describe LMAT-ML (Livermore Metagenomics Analysis Toolkit-Marker Library) which can be run with 24 GB of DRAM memory, an amount available on many clusters, or with 16 GB DRAM plus a 24 GB low cost commodity flash drive (NVRAM), a cost effective alternative for desktop or laptop users. We compared results from LMAT with five other rapid, low-memory tools for metagenome analysis for 131 Human Microbiome Project samples, and assessed discordant calls with BLAST. All the tools except LMAT-ML reported overly specific or incorrect species and strain resolution of reads that were in fact much more widely conserved across species, genera, and even families. Several of the tools misclassified reads from synthetic or vector sequence as microbial or human reads as viral. We attribute the high numbers of false positive and false negative calls to a limited reference database with inadequate representation of known diversity. Our comparisons with real world samples show that LMAT-ML is the only tool tested that classifies the majority of reads, and does so with high accuracy.

## Introduction

Recent studies show that the microbiome plays an important role in the health of humans, animals, and natural and agricultural systems. [1-4] Metagenomic sequencing of human microbiomes has already contributed to diagnosing and treating sick patients [5], and is poised to play a much larger role, provided that the technique can deliver accurate and timely analysis of multi-gigabases of unassembled reads. Metagenomic analysis typically demands substantial computing resources, either in terms of CPU or memory, or both, and run times can exceed the time for sequencing. [6] As institutions invest in sequencing infrastructure, they may not have a parallel capability to invest and maintain large compute clusters, and issues of patient privacy or data transfer bottlenecks may discourage cloud or centralized analysis. For large datasets, running BLAST analysis on Amazon’s EC2 cloud was several times more expensive than the sequencing itself, and costs of sequencing are declining faster than those of computing. [7] Rapid, sensitive, and accurate methods of taxonomic classification of the sample contents that can run on relative low price desktop machines promise a solution.

To achieve the goal of fast, accurate metagenomic analysis, various metagenome analysis software packages reduce the original sequence database to a smaller, more easily searchable marker-library containing a taxonomically informative subset. Metaphlan2 [8] matches reads to a small set of marker genes, single-copy genes present in many bacteria, or clade-specific genes to do taxonomic classification and abundance estimation without attempting to classify all reads. Kraken [9] matches the k-mers (k=31) in a read to those in a reference database. It pre-computes the lowest common ancestor (LCA) of reference sequences containing each k-mer, and applies a tree traversal algorithm to taxonomically label a read from its k-mers. The full Kraken database, however, does not fit in memory on common desktop computing systems. The MiniKraken database was created as a less memory resource intensive alternative for running on desktop systems. Clinical Pathoscope [10] aims to identify pathogens in clinical samples by alignment to NCBI bacterial and viral reference genomes and Bayesian statistical confidence estimation. GOTTCHA [11] maps reads to a pre-computed database of unique subsequences at multiple taxonomic ranks (family, genus, species, strain, etc.). SIANN [12] also maps reads to a pre-calculated database of species- and strain-specific regions of pathogens and their near neighbors. Unlike the other methods mentioned here, SIANN was designed for a specific task of rapidly assessing whether any members of a defined set of pathogens is present in a metagenomics sample. It is not a general-purpose metagenomics tool. SURPI presents another recent pathogen detection system, which uses an approach similar to Clinical Pathoscope by mapping reads to reference genomes for organism identification, but it requires more memory (60 GB) than is available on a typical desktop. [5]

LMAT uses a reference genome database that contains both draft and finished genomes from bacteria, archaea, viruses, and some eukaryotes including pathogenic protozoa. This set of reference genomes is more than 11-fold larger than any other metagenome analysis reference sequence database.[13, 14] LMAT indexes every *k*-mer (k=20) in a reference genome database with all of the sequences containing that *k*-mer. It implements a “pruning” strategy to retain only higher level taxonomic labels for *k*-mers shared by more than some pre-specified number of sequences down a taxonomic branch. It still retains multiple taxonomic nodes and genomes per *k*-mer, and thus more complete information about which sequences contain a *k*-mer than is possible with an LCA approach, which stores only a single taxonomic node per *k*-mer. This allows higher resolution (e.g. species and strain) calls when the data warrants. The full database (LMAT Grand) requires 500 GB of DRAM or flash memory so is not feasible for “desktop” or typical cluster users. LMAT-Grand’s extensive representation of genomic variation leads to labeling a large fraction of reads, which is useful for some applications (such as read binning for assembly) but is more computationally costly than may be needed for organism identification. The LMAT Marker Library (ML) reduces the RAM requirements by preselecting only the most taxonomically informative and non-overlapping (i.e. non-redundant) 20-mers for indexing, and by imposing more stringent pruning. Thus, memory requirements of LMAT-ML are reduced not by limiting the taxonomic coverage and strain resolution of the reference database, but by pre-selecting the subset of *k*-mers with the highest taxonomic information content. Moreover, the marker library approach has the potential to run at least an order of magnitude faster by correctly “ignoring” the less taxonomically informative portions of the query set. Part of the work we present here evaluated several LMAT-ML pruning levels to determine the optimal balance between memory, speed, and consistency of results compared to LMAT Grand. We also compared an LMAT-ML that contained only microbial *k*-mers to LMAT-ML+H that also contained all the human *k*-mers in LMAT Grand.

Each of the above methods makes different tradeoffs in terms of memory, speed, sensitivity, and accuracy. We chose to compare these marker library methods using data from actual microbiome data for several reasons. Previous studies with simulated datasets ensure that the simulated reads come from organisms or close relatives present in the reference database, which cannot be assumed in real samples. Constructing realistic, robust simulated metagenomic benchmarking datasets remains a fundamental challenge and will benefit from emerging community efforts to construct resources available for third party validation. While these resources develop, our goal was to compare tools, which could be run with 16 GB of RAM or less on an important target subset of metagenomics – human metagenomics using 131 Human Microbiome Project (HMP; http://hmpdacc,orq/HMASM/) samples randomly selected to span body sites and genders, and check discordant results between methods with BLAST. These samples may contain many species not included in NCBI RefSeq (http://www.ncbi.nlm.nih.gov/refseq/), or represented only as draft genomes, as well as some that have not yet been sequenced. Although we do not know ground truth, we looked at cases of disagreement between the LMAT-Grand method that queries the most comprehensive reference database and each of the marker library based methods. We examined the reads responsible for discordant species calls using BLAST [15] searches to assess if they were most likely false negatives for one method or false positives for the other. We report on the speed and accuracy of these metagenome analysis tools for HMP samples. Our evaluation considers a hardware configuration of 16 GB DRAM and a low cost commodity NVRAM in the form of a solid-state drive (SSD), which should be accessible for use with existing desktop computers. The LMAT ML databases range from approximately 13 GB to 19 GB in required storage; thus, this range covers both fitting in and exceeding the available DRAM. This range allows us to measure impact of database size on LMAT classification performance.

## Materials and Methods

### Building LMAT-ML

The LMAT reference genome database includes 1) eukaryotic sequence of fungi, protozoa, and some multicellular organisms (from organelles labeled as whole genome, e.g. mitochondria and chloroplasts); 2) draft genomes and assembled contigs from unfinished Whole Genome Shotgun (WGS) genome sequencing projects (ftp://ftp.ncbi.nih.gov/genbank/wgs/); and 3) draft and finished bacteria, virus, archaea, fungi, and protozoa genomes from a number of sequencing centers worldwide with publicly available sequence data in addition to those from NCBI^1^. An extensive collection of artificial vector sequence [17] is also included to filter out contaminating sequence in draft assemblies. The fungi and protozoa sequence data came from the Fungi and Protist Group BioProjects reported in the NCBI eukaryotes genome report (ftp://ftp.ncbi.nlm.nih.gov/genomes/GENOME_REPORTS/eukaryotes.txt), and BioProject sequences were extracted by the assembly accession.

To select the *k*-mers for inclusion in LMAT-ML, we used the procedure described in [14], using k=20 instead of 18, and adding an extra step to remove overlapping *k*-mers relative to a reference sequence (randomly selected from those containing a series of adjacent *k*-mers). Briefly, the objective was to identify a collection of *k*-mers that are uniquely associated with phylogenetically distinct sets of genomic sequence. Groups of genomes are defined by their shared *k*-mers with a minimum of 200 *k*-mers shared within a group for viral genomes and 1000 *k*-mers shared within a non-viral group. The minimum thresholds were set to maintain groups of genomes that retain some degree of phylogenetic relatedness. Any *k*-mer found in more than one group was eliminated to yield a set of *k*-mers that are uniquely associated with different levels of the taxonomy hierarchy. Additionally, *k*-mers matching RepBase18.06 [16] were eliminated. Since the resulting *k*-mer set still yielded a database with a larger memory footprint than would be practical for a desktop or laptop, *k*-mers were mapped to a randomly selected representative sequence from the genome group to remove multiple adjacent overlapping *k*-mers. Next, synthetic *k*-mers from LMAT-Grand were added to detect synthetic/vector sequences. Finally, a “+H” version of the LMAT-ML’s was created to use the human *k*-mers from LMAT-Grand to accurately classify human host sequences, while maintaining small memory requirements.

We evaluated two taxonomy pruning strategies to improve speed and reduce the memory requirements, as described in detail in [13]. Evaluating these pruning strategies is necessary to assess the possible cost in lost accuracy while improving speed. Such an evaluation is an important step in understanding the potential for LMAT-ML since the pruning strategy is a unique property of LMAT’s approach to classification, in contrast to other tools. As a baseline, when no pruning is used, indicated as “–All”, every taxonomy identifier of lowest available rank (e.g. species or strain) for a given *k*-mer is retained in the searchable database. The –Min pruning option (LMAT-ML-Min and LMAT-ML+H-Min) stores only the lowest common ancestor (LCA) for each *k*-mer, similar to the approach employed by Kraken.

The alternative pruning option tested stores a maximum of 10 taxonomy identifiers per *k*-mer (LMAT-ML and LMAT-ML+H). LMAT databases with the “+H” label include all human *k*-mers in addition to the microbial *k*-mers. In this option each *k*-mer is linked to a set of taxonomic identifiers that contain the lowest common ancestor for all sequences containing the *k*-mer and up to 9 descendent identifiers, which must retain a common rank. For example, if the LCA is of rank genus, and the k-mer is found in nine or fewer distinct species, then all distinct species identifiers would be retained. If the k-mer is found in more than nine species (but only one genus) then only the genus identifier would be retained.

The LMAT database uses a two-level index data structure described in detail in [19] to improve the efficiency of *k*-mer search. A *k*-mer is represented by two non-overlapping bit vectors, with a 20-mer represented by 40 bits and the first N bits stored in the first level of the index and the second 20-N lower order bits stored in the second level. The choice of N was optimized for use with the ML. A split of N=25 bits was selected, which reduced the size of the ML database by 1.5 GB, compared with previous settings developed for use with the full database. A new extension to the LMAT software was added (v 1.2.5), enabling the two-level split to be specified for the target database. This feature allows each database to be uniquely tuned for efficient use of space adjusting for the size of the database. The split parameter was adjusted from the previous setting used for the larger database (LMAT-Grand) to deploy the smaller marker databases on a low cost SSD device.

### Marker library comparisons

We used a set of 131 HMP samples randomly chosen to span all body sites for both genders (Additional file 1: 131HMP_Samples.xlsx), with preprocessing to trim non-biological portions of reads (adaptors), trim or replace low quality bases (Q<10) with N, and combined paired end sequences as described in [14]. We compared results from Clinical Pathoscope v1.0.3, Metaphlan2 (db_v20), GOTTCHA (downloaded Sept. 2, 2014), MiniKraken (kraken-0.10.4-beta), SIANN (v1.12), the LMAT-ML+H, and the LMAT Grand database. Each method was run with default parameters. Clinical Pathoscope target databases were bacteria and virus, and host filtration database was human. GOTTCHA was run against the GOTTCHA_BACTERIA_c3514_k24_u24_xHUMAN3x.species and GOTTCHA_VIRUSES_c3498_k85_u24_xHUMAN3x.species databases, microbial databases in which 24-mers matching the human reference genome had been removed, and results were combined for bacteria and viruses. Results from all LMAT-MLs with no or moderate pruning were nearly identical in terms of consistency with the LMAT-Grand database, while results were slightly worse for LMAT-ML-Min without human k-mers (Figure S1), so for all the marker method comparisons we used LMAT-ML+H. We uniformly applied the default cutoff values of 0.5 for LMAT-Grand and 0.2 for LMAT-ML+H. We required a minimum of 100 reads per species to call that species as present for each of the methods, and observed that results were similar across thresholds of 50-2000 reads (results not shown). Reads for methods that reported calls as percentages of mapped or total reads were converted to absolute numbers of reads. For Metaphlan2, since the relative abundance was not straightforward to convert to read counts, we used the percent abundance multiplied by the total number of reads mapped as a proxy. We identified the NCBI species-level taxonomy id and NCBI species name for all species and strain calls by each method, a nontrivial process since some methods report non-standard species names and no taxonomy identifiers, and use deprecated GI numbers, which are not in the current NCBI databases.

For each method, the 10 most commonly called species detected by that method and not by LMAT-Grand were identified, and the reads identified by that method as belonging to that species were extracted based on parsing the bowtie2 or SAM output, identifying the taxonomy by GI number, NCBI gene ID (Metaphlan2), or as a last resort by organism name, using NCBI tables^2^. We acknowledge that we may have missed extracting some reads which a given method used for classification, since these methods do not describe standardized procedures for read extraction and call validation. For Metaphlan2 in particular, we were unable to extract as many reads as we expected based on reported percentage abundances. Reads were combined into a single file per species for each method. These reads were then compared using BLAST (blastn -evalue 0.0001 -max_target_seqs 5) to our comprehensive database of all bacterial and viral sequences, and the output pruned to show only the matches to each read with the lowest e-values, allowing multiple matches per read with the same lowest e-value. These reads were also compared using BLAST to a comprehensive database of vectors and other artificial sequences, containing the sequences in UniVec as well as Illumina adaptors and a number of commercially available vector sequences [17]. The reads were also compared using LMAT with the Grand database, and the taxonomic call with highest read count was reported.

The 10 most common species calls made by LMAT-Grand and not another method were gathered for each method, ignoring Homo sapiens and LMAT-Grand classification for synthetic constructs, which LMAT-Grand reports as a “species". The top 10 list was identical for Clinical Pathoscope, GOTTCHA, and MiniKraken, which all use NCBI RefSeq to build their reference databases. Reads for these species with LMAT-Grand scores of at least 1 were extracted and compared using BLAST against our comprehensive viral/bacterial genome database, and in one case where a protozoa was uniquely detected, against our protozoa database.

### Results

LMAT databases encode 11-147 times more sequence data representing approximately 2-4 times more species than other methods (Table 1). The memory (DRAM or DRAM+NVRAM) of LMAT-ML is also higher than for other methods, although still feasible for a desktop, particularly when supplemented with low cost NVRAM.

**Table 1:**
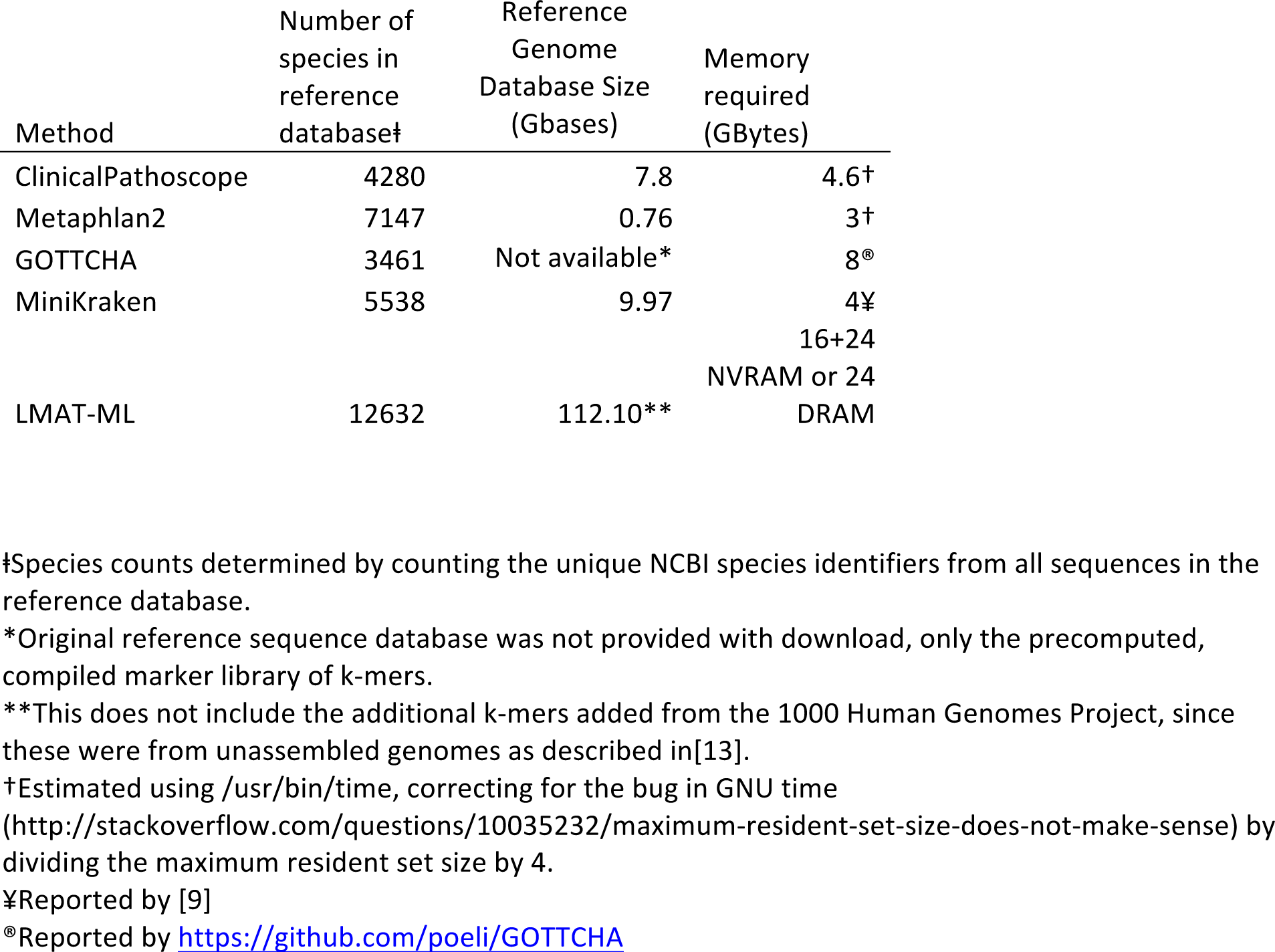
Reference genome database size and number of species represented by each method.

### Runtime performance is determined by database size

Table 2 shows the size of the different ML databases reflecting the different pruning strategies (shown as 1, 10, and All in third column of the table). Although there was little difference in taxonomic classifications between 4 of the 5 LMAT-ML (Figure S1), the choice of pruning level affects memory requirements. For comparison, LMAT-MLs include approximately 4.4-4.7 times more *k*-mers than MiniKraken but require 3-4.7 times more memory. However, LMAT-ML-Min maintains a lower *k*-mer per byte ratio than MiniKraken (7.6 versus 11.2), which is likely explained by LMAT’s use of a smaller *k* (20 versus 32 for MiniKraken).

**Table 2:** Database sizes and k-mer counts (for LMAT) given configurations that vary the presence of human reference k-mers and the maximum count of taxonomy ids listed per k-mer. *For comparison we include an estimate of MiniKraken’s k-mer count based on 12 bytes per *k*-mer as provided by the authors.[9]

| Method | Contains human k-mers? | Maximum taxonomy ids stored per k-mer | size (GB) | K-mers |
| --- | --- | --- | --- | --- |
| MiniKraken | - | 1 | 4.0 | 357,913,941* |
| LMAT-ML-Min | no | 1 | 12.1 | 1,586,405,299 |
| LMAT-ML | no | 10 | 15.9 | 1,586,405,299 |
| LMAT-ML+H | yes | 10 | 16.8 | 1,697,066,355 |
| LMAT-ML-All | no | All | 18.1 | 1,586,405,299 |
| LMAT-ML+H-All | yes | All | 18.9 | 1,697,066,355 |

**Figure 1:**
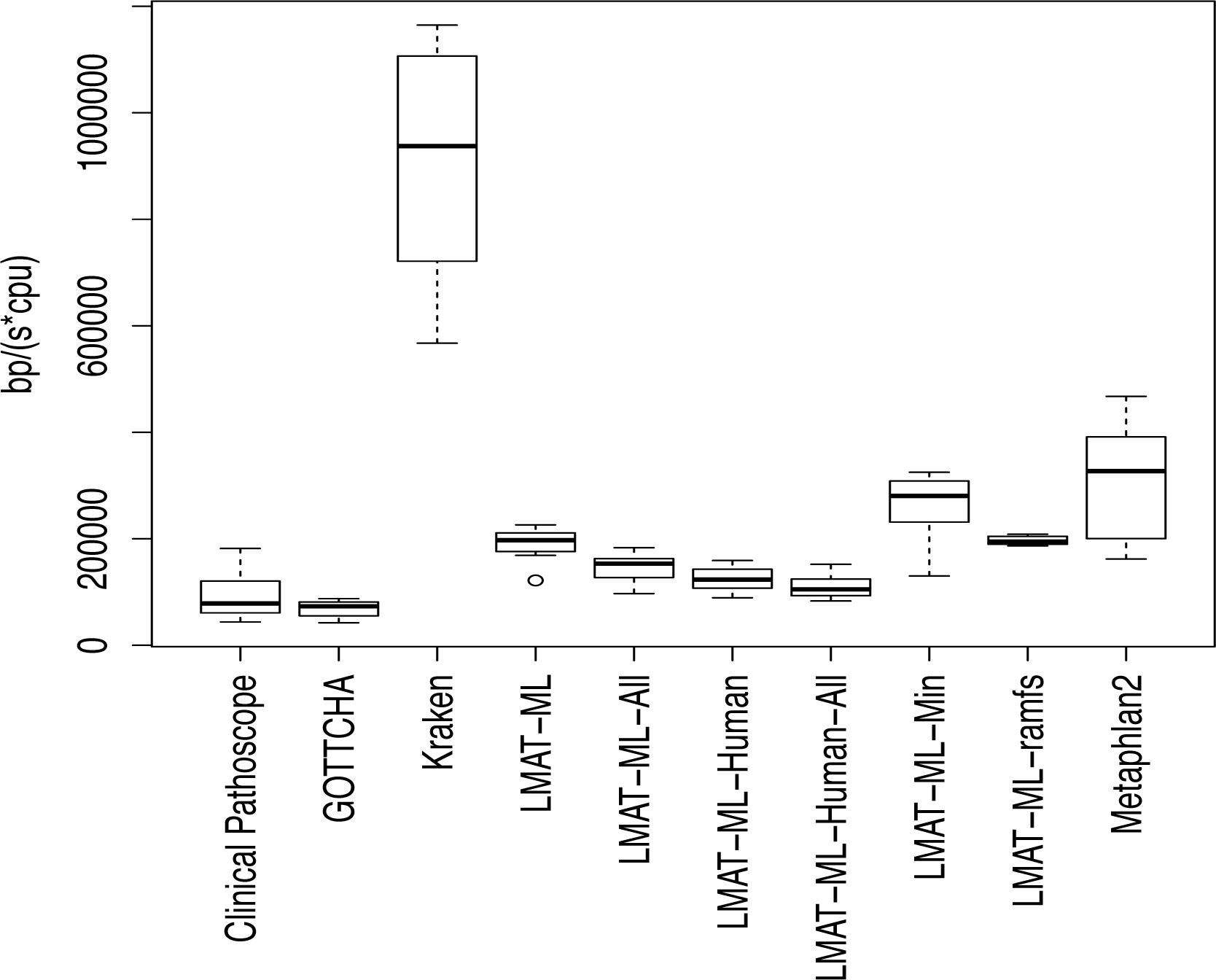
Processing rate for LMAT using 5 database configurations and four additional processing methods, reported per CPU. 5 of the 6 LMAT-ML runs use SATA II SSD for storage. LMAT-ML-ramfs, the LMAT database is stored in main memory on a “ramdisk” (linux “ramfs”) file system. In this and the following figures, MiniKraken is labeled simply as ‘Kraken’.

Figure 1 shows processing rates for LMAT compared with the four classification tools. Each box plot bar encompasses the results from 8 HMP sample runs. For the LMAT runs we have configured five marker libraries, and query them using 16 GB of DRAM and a SATA II Ocz 500 GB solid-state drive. To compare performance when storing the database completely in DRAM memory, we show the performance of the LMAT-ML database on a compute platform with 24 GB of DRAM, where the database index can reside in main memory without paging from the external storage device using a ramdisk (linux ramfs), as indicated by label LMAT-ML-ramfs. From these results we make several observations.

1. (1) MiniKraken has approximately 5 times faster performance than our baseline LMAT (at most 10 taxonomy identifiers per *k*-mer). Its database size is considerably smaller at 4 GB in contrast to the LMAT databases (Table 2); thus, it should have greater CPU cache hit rates on average, which can have a profound impact on performance.
2. (2) LMAT-ML-Min (singe taxonomy identifier per *k*-mer) shows faster performance than our ramdisk run (LMAT-ML-ramfs). This identifies the increase in cost of processing more taxonomy identifiers.
3. (3) LMAT-ML (SSD configuration) is within 4% of LMAT-ML-ramdisk. While we estimate that as much as 15% of the database would not be available in buffer cache at any moment, this result indicates that this ratio of database size to total DRAM does not create a substantial burden on the system’s caching mechanism.
4. (4) The LMAT-ML+H and both LMAT-ML-All databases show slower performance. In all cases, the larger database size means less of it can fit in buffer cache, thus requiring more NVRAM access operations. The addition of human appears to have a greater impact on performance than the increase in taxonomy identifiers per *k*-mer indicating that the number of distinct *k*-mers stored in the index necessitates more NVRAM access operations.
5. (5) Metaphlan2 is faster than all LMAT-ML instances, but LMAT-ML-Min is within 20%.
6. (6) All LMAT configurations outperform GOTTCHA and Clinical Pathoscope in terms of speed, although Clinical Pathoscope is close to the LMAT configurations that contain human reference information.

### Comparing the organisms detected by each method

SIANN found none of the organisms in its database for any of the HMP samples. We manually searched for several pathogens that were included in their database and which were detected by both Metaphlan2 and LMAT-Grand but not SIANN, and these included *Clostridium symbiosum, Burkholderia cenocepacia, Staphylococcus epidermidis, S. caprae, S. hominis, S. lugdensis*, and *S.aureus*. We ran it on several samples with known spiked pathogen concentrations [18] to verify that we were running the software correctly. In a sample with 10,000 genome equivalents (GE) of *Bacillus anthracis, Burkholderia pseudomallei, Francisella tularensis*, and *Yersinia pestis* and an unknown concentration of *Brucella* spiked into a background of human DNA and sequenced on Ion Torrent, SIANN detected the 5 spiked pathogen species with high confidence but was unable to make a correct strain call. It also made 2 false positive species calls for organisms that were not present *(Francisella novicida* and *Francisella cf*). At 100 GE spike in, only an incorrect strain of *F.tularensis* was detected with low confidence, and none of the other pathogens present were detected. LMAT Grand and LMAT-ML+H detected all 5 pathogens at 10,000 and 100 GE. LMAT called the vast majority of reads at the species level (except for Brucella genus) indicating these regions are conserved across multiple genomes. LMAT with 10,000 GE correctly called *Y. pestis* Harbin as the top strain, and in the other cases the top strain was among the top 3 genomes with the most genome-specific reads. We did not investigate SIANN further, as it was not designed to be a general purpose metagenomics analysis tool.

An important component of identifying the organisms present in a metagenome is measuring the number of reads left unclassified. Even as microbial diversity in genomic databases continues to grow, the potential for novel genes or organisms to remain ‘hidden’ in the sample remains high. Thus, the ability to reduce the number of reads that must be considered for assembly or other more in depth analysis through time consuming sensitive protein searches with BLAST or profile Hidden Markov Models must be considered when evaluating the completeness of a method’s taxonomic profiling. LMAT-Grand classified on average 83% of the reads per sample, followed by LMAT-ML+H (63%), MiniKraken (35%), Clinical Pathoscope (29%), GOTTCHA (14%), and Metaphlan2 (5%) (Figure 2A). LMAT-Grand detected an average of 178 species/sample, LMAT-ML+H 154 species/sample, MiniKraken 108 species/sample, Clinical Pathoscope 67 species/sample, Metaphlan2 42 species/sample, and GOTTCHA 27 species/sample (Figure 2B).

**Figure 2A:**
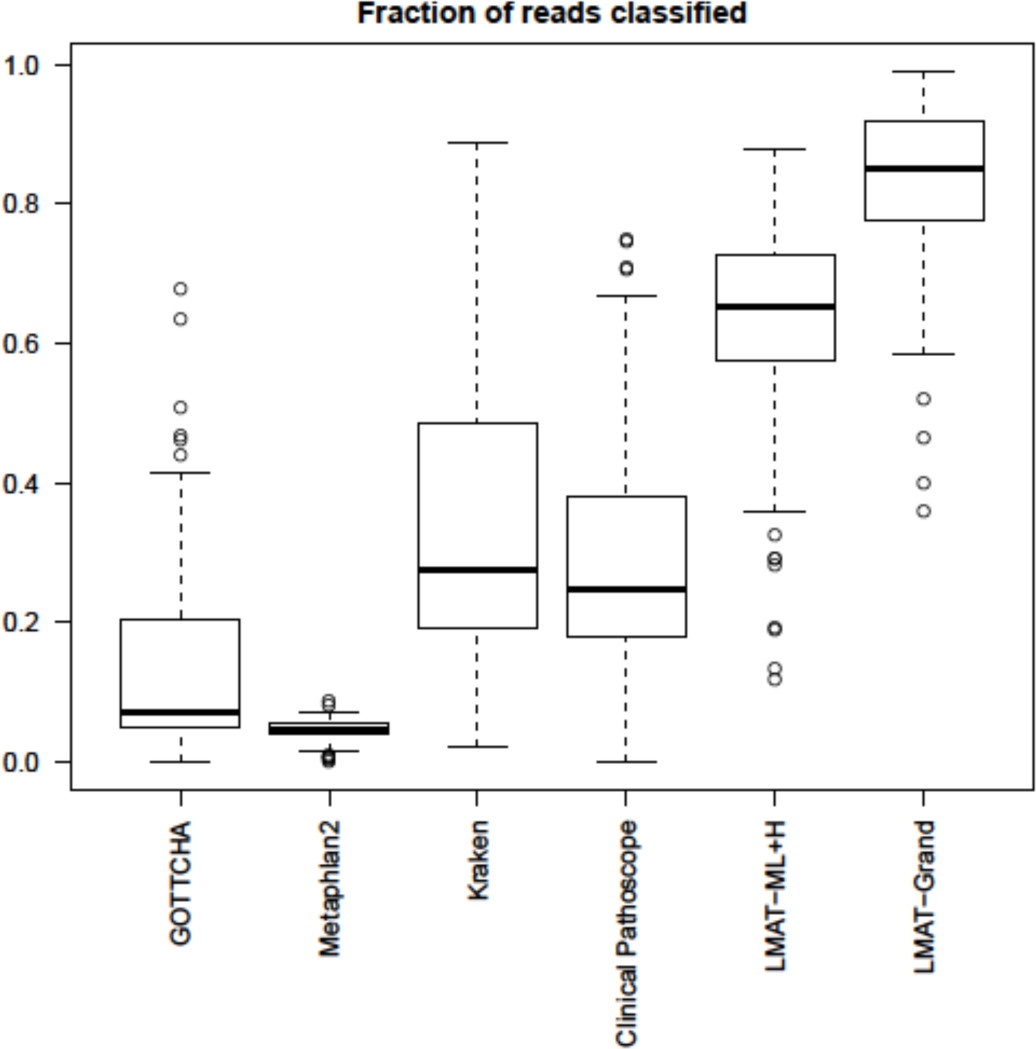
Boxplots showing the fraction of reads that each method classified from the 131 HMP samples.

**Figure 2B:**
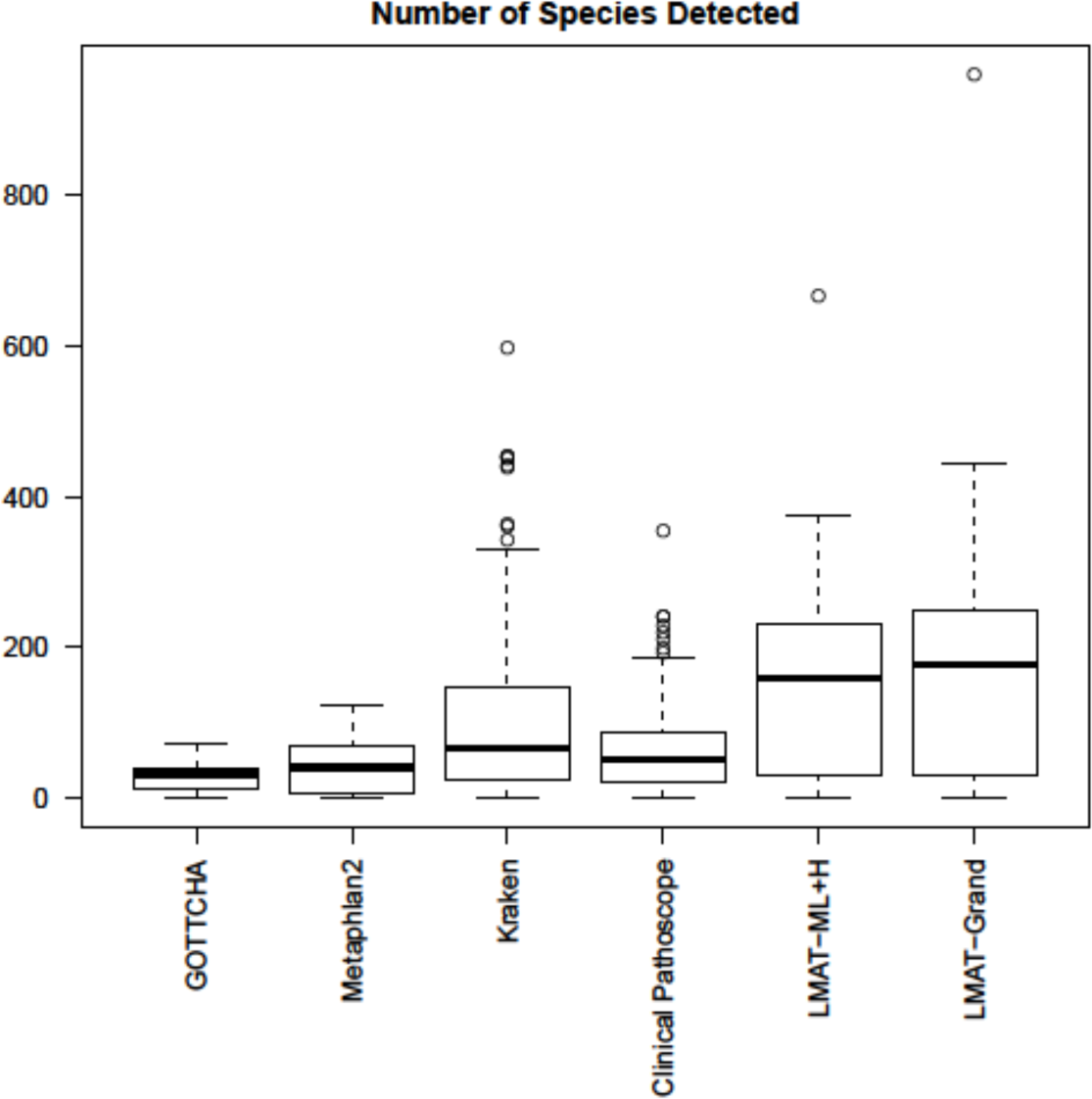
Boxplots of the number of species that each method classified from each of the 131 HMP samples. LMAT Grand detected on average 11% more species than LMAT-ML+H and 57% more species than MiniKraken, the next highest method. BLAST results suggest a large number of MiniKraken calls are false positives.

Taxonomy calls made using LMAT-Grand do not represent a complete ground truth of organisms present in a sample. Since the database draws on the most complete collection of sequenced genomes among databases considered it is useful to measure the relative concordance among the different methods relative to LMAT-Grand. The overlap between each method and LMAT-Grand differed substantially as illustrated in Figure 3, summing species calls in agreement or in disagreement with those of LMAT Grand across the 131 samples. MiniKraken and Clinical Pathoscope showed relatively little overlap with LMAT-Grand species calls, GOTTCHA and Metaphlan2 overlapped by the majority of their species calls but only covered a small fraction of the LMAT Grand calls, and LMAT-ML+H and LMAT Grand agreed almost entirely.

**Figure 3:**
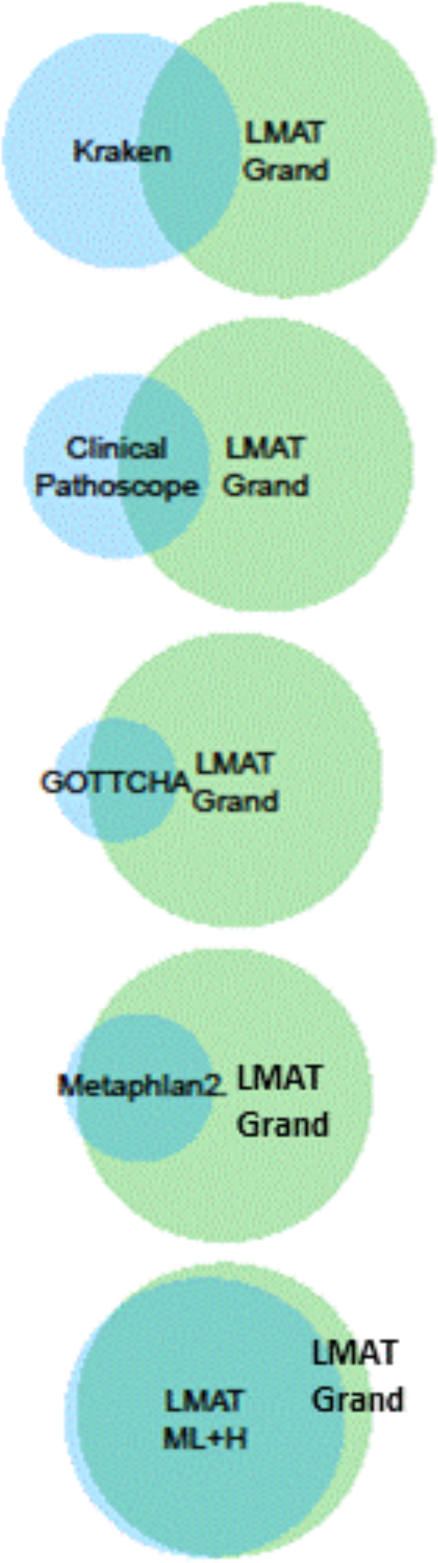
Venn diagrams illustrating the total number and overlap of species calls for the 131 HMP samples by each method with LMAT Grand, in order of increasing overlap.

The calls by GOTTCHA, MiniKraken, Metaphlan2, and Clinical Pathoscope share higher similarity with those of LMAT-Grand at the genus level than at the species level, although neither species nor genus agreement is very high (Figure 4). To consider the possibility that the LMAT-Grand output was error prone, a detailed analysis of the discordant calls were examined using BLAST alignments against a comprehensive microbial database.

**Figure 4:**
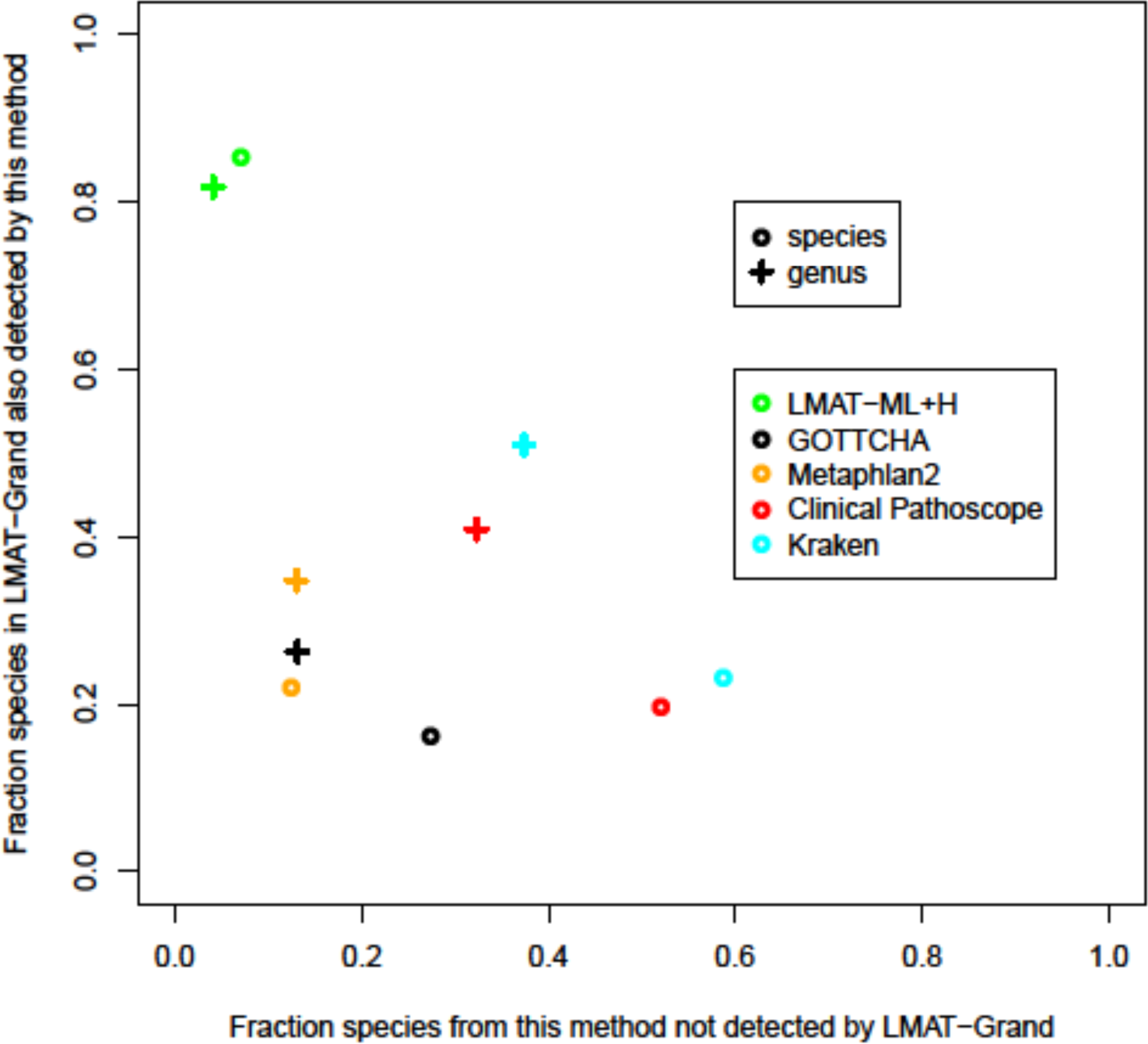
The fraction of species (circles) or genera (crosses) detected by LMAT Grand that were also detected by a given method versus the fraction of species detected by a given method that were not detected by LMAT Grand, averaged across the 131 HMP samples. If the calls by LMAT Grand are correct as indicated by BLAST results (see below), then this is akin to sensitivity versus false discovery rate.

### Calls made by other methods that were not supported by LMAT Grand

All the marker library methods except LMAT-ML detected a number of species in each sample that LMAT-Grand did not (Figures 3 and 4). Reads were extracted from a given method’s mapping results for the 10 most commonly occurring species calls that differed from LMAT Grand. Figure 5 shows the sum of these extracted reads across samples and species, totaling between 10,000 to greater than 1M reads for non-LMAT methods. With orders of magnitude fewer reads for LMAT-ML+H potential false positives, manual inspection revealed that the difference between LMAT-ML+H and LMAT-Grand lay in minor quantitative differences close to the threshold read count for calling a species as present rather than qualitatively different calls. BLAST searches with these reads against the comprehensive microbial genome database provided evidence in support or contradiction of the potential false positives, summarized in Figure 6 and described in detail for each method.

**Figure 5:**
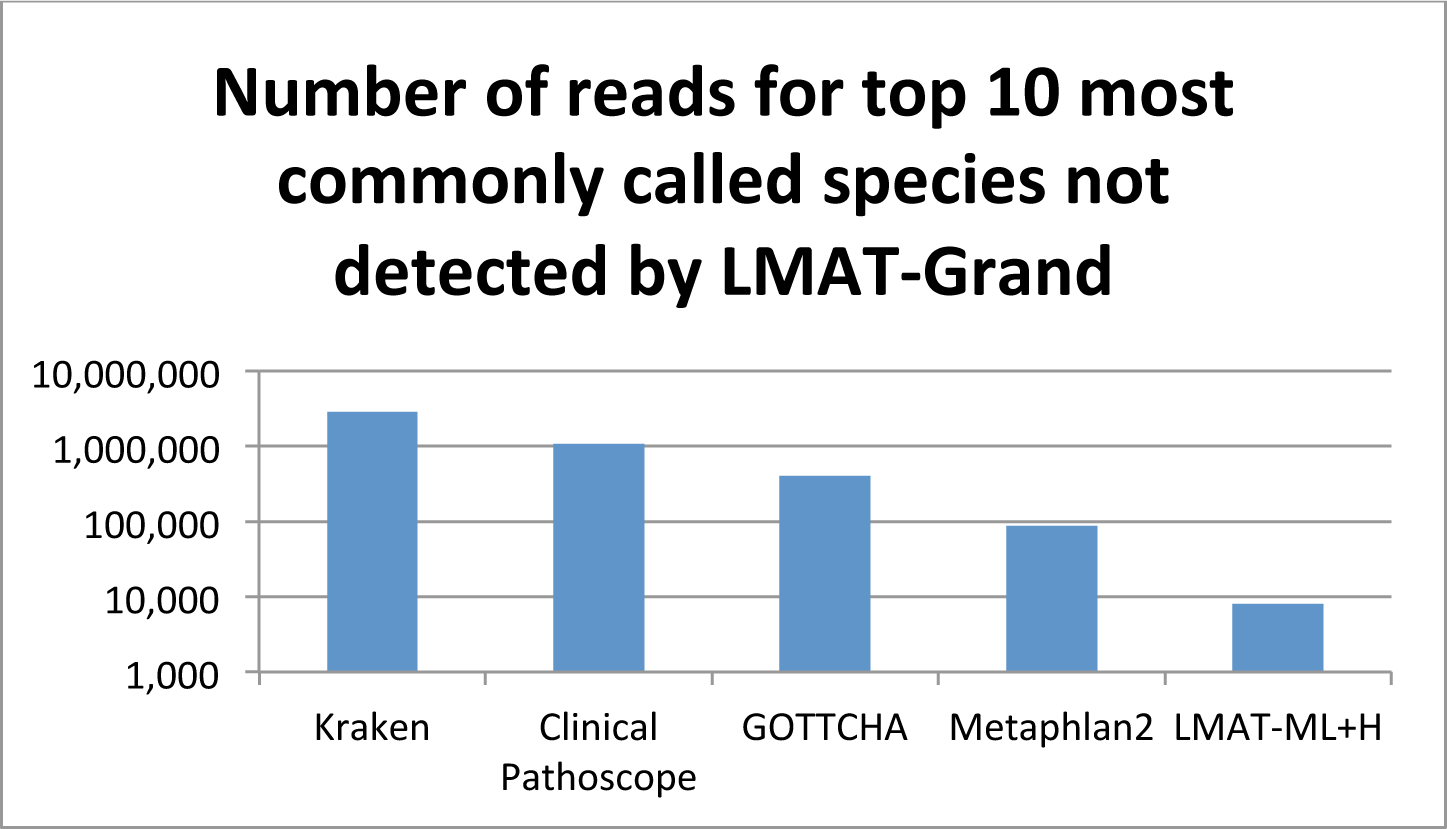
Number of candidate false positive reads totaled across samples for the top 10 most common species calls that were not detected by LMAT Grand.

**Figure 6:**
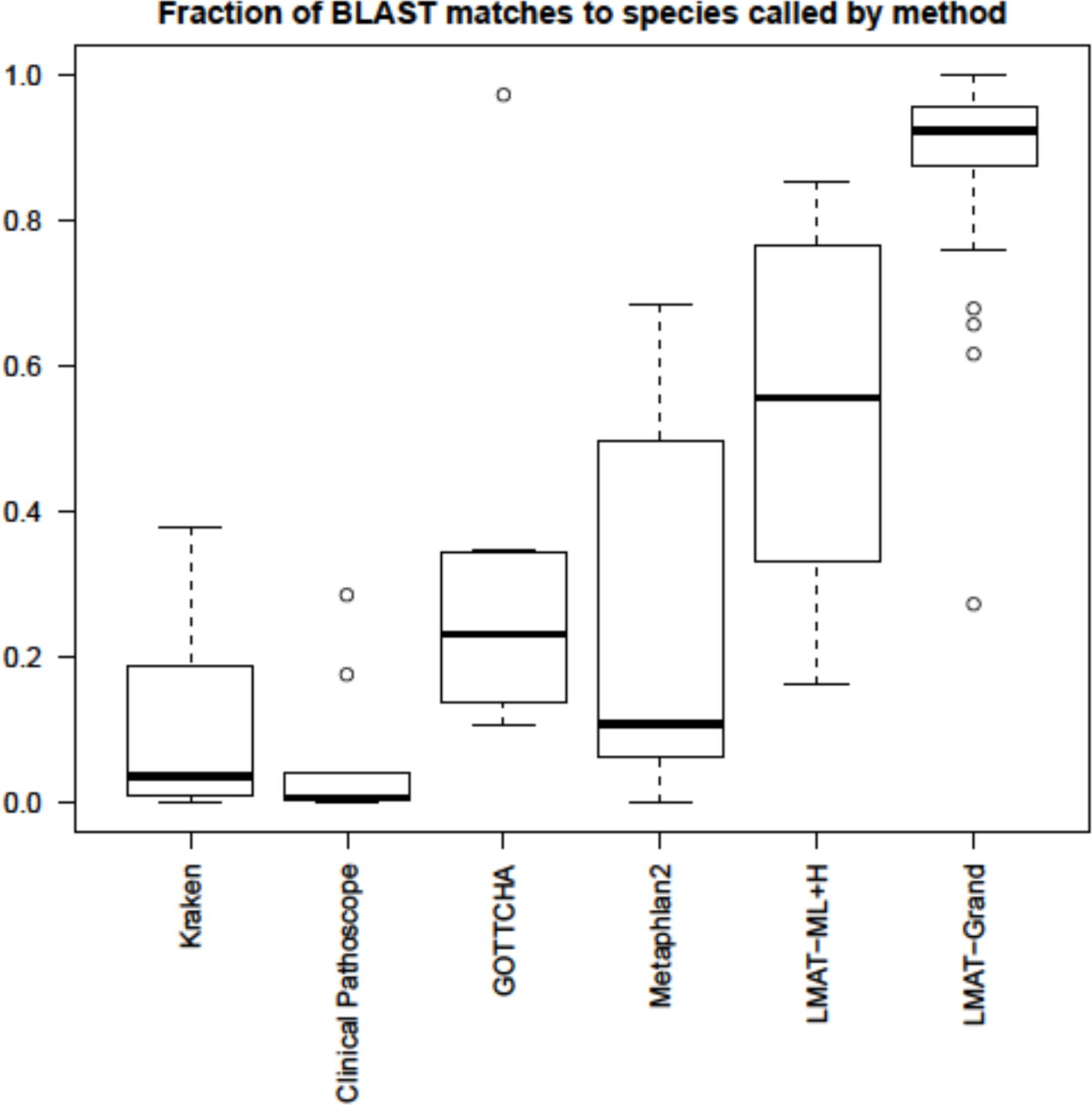
Fraction of best BLAST matches to species called by the indicated method that were discordant with species calls by LMAT-Grand.

These were for the most commonly called species by the indicated method that were not detected by LMAT-Grand in those same samples. In all non-LMAT methods, only a minority of BLAST hits to a comprehensive sequence database supported the species called by that method, suggesting that these were incorrect species calls. Calls by LMAT-ML+H had a majority of matches to the correct species, on average, but nearly as many matches to other species in the genus, supporting genus level calls for some of these reads. BLAST results provided strong support for LMAT- Grand calls that were not detected by other methods. A more detailed analysis of the discordant calls for each method is now given in the following sections.

### MiniKraken unsupported calls

MiniKraken made a number of unsupported calls that were repeatedly seen across half or more of the samples. For the most common MiniKraken species calls that were not consistent with LMAT species calls, 7 of the 10 had 5% or fewer BLAST hits to the species identified by MiniKraken. The other 3 still had a minority with 19-38% of BLAST matches to the MiniKraken species, and the most common LMAT classification for these reads was at the genus level or above. The unsupported MiniKraken species call with the bulk of reads, over 2 million, had 99.5% of reads matching the database of synthetic sequences. These BLAST results indicate that MiniKraken calls were either incorrect and/or overly specific, and failed to correct for contamination of reference sequences with vector and other artificial sequence contaminants (Supplementary Table S2).

### Clinical Pathoscope unsupported calls

Clinical Pathoscope commonly called a number of bacteria in nearly half the samples which were not supported by LMAT-Grand calls. Of reads mapped to organisms called by Clinical Pathoscope but not LMAT-Grand, most had fewer than 1% of BLAST hits to this organism, suggesting that organisms with the highest similarity to these reads were not in the ClinicalPathoscope database, so the reads were mis-attributed to other species with homology (Supplementary Table S3). There were two exceptions for which more than 10% of the lowest e-value BLAST matches were to the organism called by Clinical Pathoscope: *Propionibacterium acnes* with 28%, and *Bacteroides vulgatus* with 17%, although these are still a minority of BLAST matches. LMAT-Grand classified most of those *P. acnes* reads as genus, *Propionibacterium*, order, *Actinomycetales*, class, *Actinobacteria*, and cellular organisms. LMAT-Grand called the majority of those *B. vulgatus* reads at the *Bacteroides* genus level, in addition to some thousands of reads to other genera, order *Bacteroidales*, phylum *Bacteroidetes*, and superkingdom Bacteria. The fact that a minority of BLAST hits are to the organism identified by Clinical Pathoscope suggest that calls by Clinical Pathoscope may be overly specific, with equivalent similarity to many species. In addition, up to 19% of the reads for some of these unsupported Clinical Pathoscope calls had BLAST matches to artificial sequences, indicating, this method fails to adequately control for contamination of the reference database by artificial sequences such as vectors and adaptors.

### GOTTCHA unsupported calls

Phage dominated the calls unique to GOTTCHA. All the reads which GOTTCHA labeled as *Enterobacteria* phage lambda matched our vector and other synthetic sequences database, and LMAT labeled almost all these reads as synthetic constructs or root (Supplementary Table S4). 23% of the reads that GOTTCHA called as *Enterobacteria* phage phiX174 sensu lato had BLAST matches to our vector database, and LMAT labeled them as root, superkindgom Bacteria, synthetic sequences, or cellular organisms. Elements of these phage are used for genetic engineering and as controls in certain Illumina sequencing protocols, so we include them in the vector/synthetic database, so this result is not surprising. While the other calls did not have many BLAST matches to artificial sequences, only a minority of the top BLAST hits matched the GOTTCHA call since these reads were widely conserved across species or higher ranks, and LMAT identified these reads largely as root or genus level calls.

### Metaphlan2 unsupported calls

There were 4 organisms commonly called by Metaphlan2 that were unsupported by LMAT-Grand calls: Dasheen Mosaic virus, *Streptococcus sp. GMD4S, Catonella morbi*, and *Abiotrophia defectiva*, plus some other unsupported calls that occurred in just a few samples (Supplementary Table S5). In contrast to Clinical Pathoscope, Metaphlan2 unique calls had very few matches to artificial sequence. Reads that Metaphlan2 uniquely labeled as Dasheen mosaic virus and Vicia cryptic virus had no best BLAST matches to these viruses, prompting us to double check that these viruses were indeed present in our BLAST database. LMAT calls for these reads were predominantly to *Homo sapiens*, plus small numbers to high level classifications such as taxonomy root node, cellular organisms, and various bacterial genera. For some of the organisms, 48%-68% of the BLAST matches were to the correct organism, but there were as many matches of the same quality to other organisms, suggesting overly specific Metaphlan2 calls to reads conserved at the genus or phylum level. Supporting this observation, LMAT calls for those particular reads were overwhelmingly at the phylum and genus levels, and even some to the kingdom Bacteria and to cellular organisms. For other Metaphlan2 species calls, LMAT Grand classified more of those reads to several near neighbor species in the same genus, with only a small subset of those reads classified as the same species as Metaphlan2 but not enough to reach the minimum call threshold of 100 reads in those samples.

### LMAT-ML+H unsupported calls

Unsupported calls were less common and less consistent across multiple samples for LMAT-ML+H than for the other methods, and involved fewer reads (Figure 5). When we investigated LMAT-ML+H differences from the Grand database, most of the differences were in samples where the number of reads called hovered around the 100 read threshold, so that species called by LMAT-ML+H were also observed in the Grand results but at abundances just below 100 reads (Table 3). LMAT-Grand calls for the extracted species reads from the LMAT-ML+H runs always included some species calls to the species detected by LMAT-ML+H for those reads, as well as a larger number at a higher taxonomy level. Manual inspection of many examples showed that both LMAT-ML+H and LMAT-Grand always called multiple species in the genus, although the distribution of reads counts among those species differed. The average score per species was usually lower for LMAT-ML+H than Grand. LMAT calculates a log odds read score from the number of *k*-mer matches in a read relative to a null model simulated with random sequences for each database, adjusting for GC content and read length of the null model to match that of the read.[14] The LMAT-ML+H database has many fewer *k*-mers than the Grand database, resulting in lower average scores. Low scores indicate an organism has best similarity to that taxonomic call, but does not perfectly match the reference sequence, suggesting a novel variant. The calls to *Tetrahymena thermophila* had average read scores of just above 0.2 for both LMAT-ML+H and Grand, and so were below the threshold for Grand but not LMAT-ML+H. These reads had BLAST matches only to *Plasmodium yoelii* and *Tetrahymena thermophila*, and were very repetitive and AT rich. Classification of such low complexity reads is a challenge, especially considering the difficulty of assembling reference genomes for protozoa around such regions, with the potential for misassembly and contamination with host and other eukaryotic DNA. In summary, differences between LMAT Grand and LMAT-ML+H were more likely for organisms with low numbers of reads or low scores, i.e. those with low abundance or low similarity to that genome in the database.

**Table 3:**
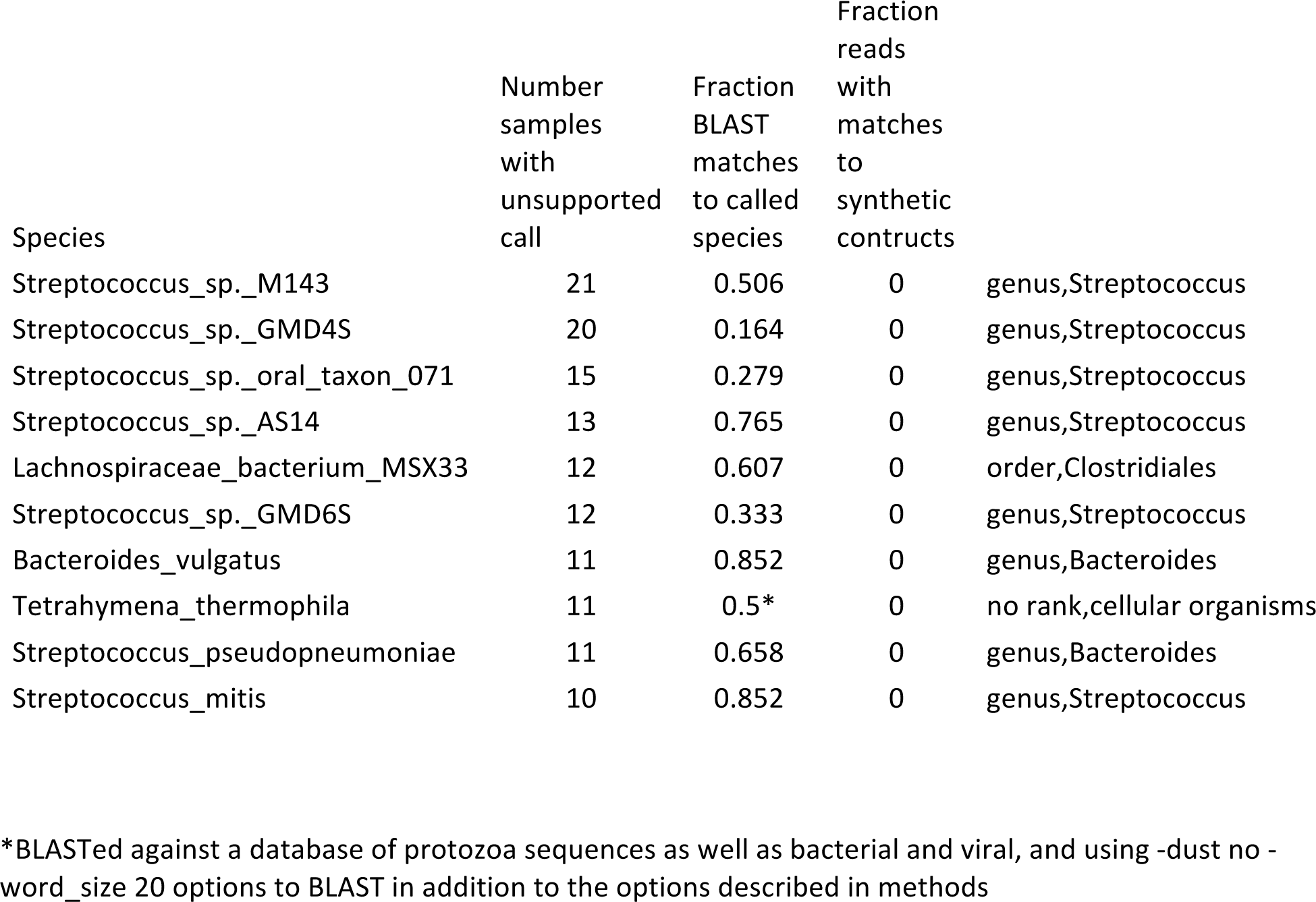
BLAST analysis of unsupported calls by LMAT-ML

| Species | Number<br>samples<br>with<br>unsupported<br>call | Fraction<br>BLAST<br>matches<br>to called<br>species | Fraction<br>reads<br>with<br>matches<br>to<br>synthetic<br>constructs |  |
| --- | --- | --- | --- | --- |
| Streptococcus_sp._M143 | 21 | 0.506 | 0 | genus,Streptococcus |
| Streptococcus_sp._GMD4S | 20 | 0.164 | 0 | genus,Streptococcus |
| Streptococcus_sp._oral_taxon_071 | 15 | 0.279 | 0 | genus,Streptococcus |
| Streptococcus_sp._AS14 | 13 | 0.765 | 0 | genus,Streptococcus |
| Lachnospiraceae_bacterium_MSX33 | 12 | 0.607 | 0 | order,Clostridiales |
| Streptococcus_sp._GMD6S | 12 | 0.333 | 0 | genus,Streptococcus |
| Bacteroides_vulgatus | 11 | 0.852 | 0 | genus,Bacteroides |
| Tetrahymena_thermophila | 11 | 0.5* | 0 | no rank,cellular organisms |
| Streptococcus_pseudopneumoniae | 11 | 0.658 | 0 | genus,Bacteroides |
| Streptococcus_mitis | 10 | 0.852 | 0 | genus,Streptococcus |

\*BLASTed against a database of protozoa sequences as well as bacterial and viral, and using -dust no -word\_size 20 options to BLAST in addition to the options described in methods

### Calls made by LMAT-Grand that were not detected by other methods

LMAT-Grand calls had strong BLAST support: the LMAT-identified species dominated the BLAST matches. This supports the notion that these are false negatives by GOTTCHA, MiniKraken, Clinical Pathoscope, and Metaphlan2, all of which use substantially smaller reference databases than LMAT-Grand (Table 4). These were not isolated false negatives, as most occurred in over half the HMP samples we compared. In contrast, for LMAT-ML+H, even the most common false negatives occurred in less than a third of the samples. For the LMAT-ML+H comparisons, in every case we examined, the species was actually detected by the LMAT-ML+H but it fell under the 100 read count threshold, so it was a matter of slight quantitative differences near the threshold rather than qualitative differences, the same result as discussed above for the LMAT-ML+H unique calls. This suggests that when using LMAT-ML+H, adjustments should be made to account for lower sensitivity of LMAT-ML+H than LMAT Grand. LMAT-ML+H uses the identical database of reference genomes as LMAT Grand, downselecting to a fraction of the most taxonomically informative *k*-mers and removing redundancy by eliminating many of the overlapping *k*-mers, so it is not surprising that differences between the two databases are small.

**Table 4:**
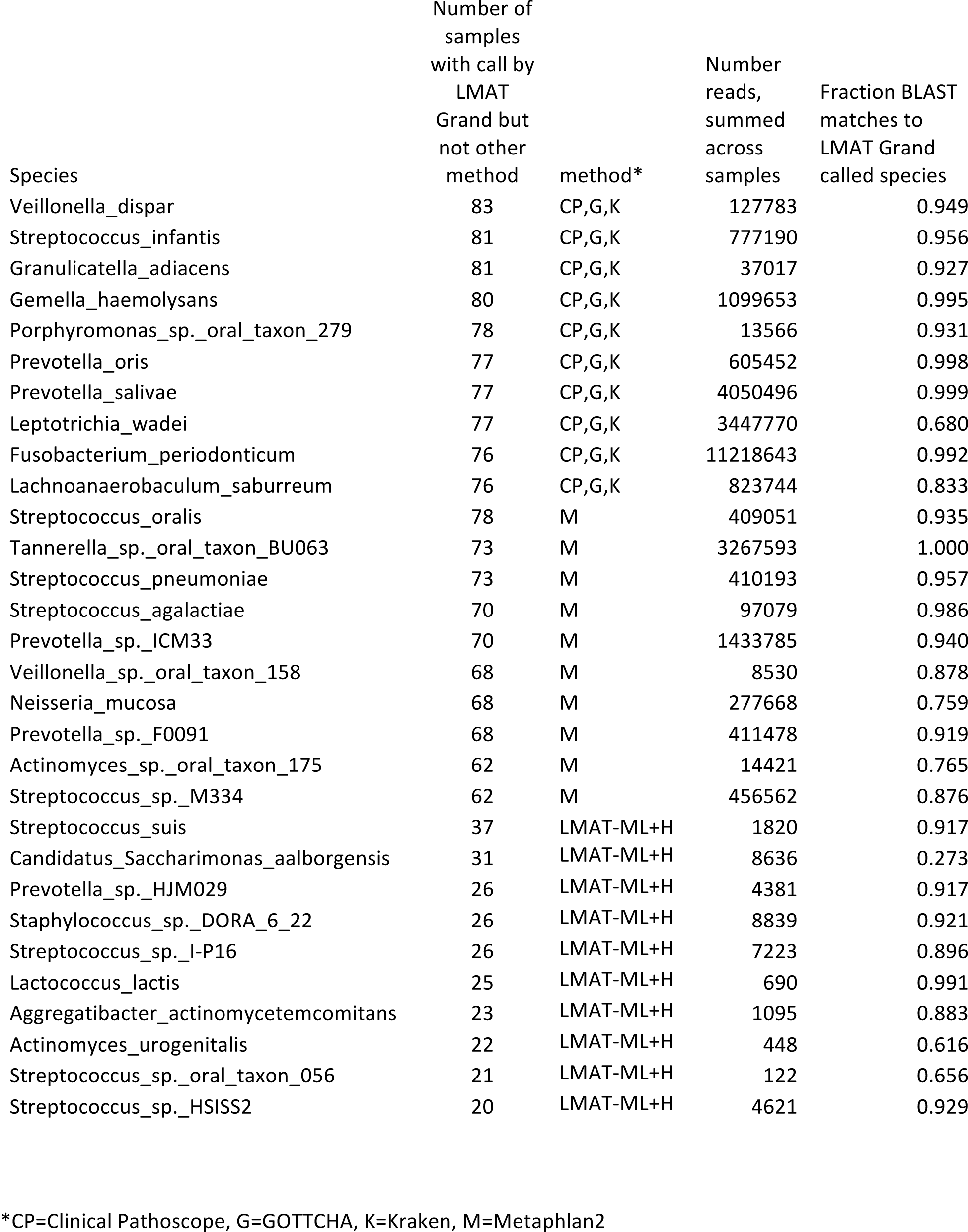
BLAST analysis of calls made by LMAT Grand but not other methods.

| Species | Number of<br>samples<br>with call by<br>LMAT<br>Grand but<br>not other<br>method | method* | Number<br>reads,<br>summed<br>across<br>samples | Fraction BLAST<br>matches to<br>LMAT Grand<br>called species |
| --- | --- | --- | --- | --- |
| Veillonella_dispar | 83 | CP,G,K | 127783 | 0.949 |
| Streptococcus_infantis | 81 | CP,G,K | 777190 | 0.956 |
| Granulicatella_adiacens | 81 | CP,G,K | 37017 | 0.927 |
| Gemella_haemolysans | 80 | CP,G,K | 1099653 | 0.995 |
| Porphyromonas_sp._oral_taxon_279 | 78 | CP,G,K | 13566 | 0.931 |
| Prevotella_oris | 77 | CP,G,K | 605452 | 0.998 |
| Prevotella_salivae | 77 | CP,G,K | 4050496 | 0.999 |
| Leptotrichia_wadei | 77 | CP,G,K | 3447770 | 0.680 |
| Fusobacterium_periodonticum | 76 | CP,G,K | 11218643 | 0.992 |
| Lachnoanaerobaculum_saburreum | 76 | CP,G,K | 823744 | 0.833 |
| Streptococcus_oralis | 78 | M | 409051 | 0.935 |
| Tannerella_sp._oral_taxon_BU063 | 73 | M | 3267593 | 1.000 |
| Streptococcus_pneumoniae | 73 | M | 410193 | 0.957 |
| Streptococcus_agalactiae | 70 | M | 97079 | 0.986 |
| Prevotella_sp._ICM33 | 70 | M | 1433785 | 0.940 |
| Veillonella_sp._oral_taxon_158 | 68 | M | 8530 | 0.878 |
| Neisseria_mucosa | 68 | M | 277668 | 0.759 |
| Prevotella_sp._F0091 | 68 | M | 411478 | 0.919 |
| Actinomyces_sp._oral_taxon_175 | 62 | M | 14421 | 0.765 |
| Streptococcus_sp._M334 | 62 | M | 456562 | 0.876 |
| Streptococcus_suis | 37 | LMAT-ML+H | 1820 | 0.917 |
| Candidatus_Saccharimonas_aalborgensis | 31 | LMAT-ML+H | 8636 | 0.273 |
| Prevotella_sp._HJM029 | 26 | LMAT-ML+H | 4381 | 0.917 |
| Staphylococcus_sp._DORA_6_22 | 26 | LMAT-ML+H | 8839 | 0.921 |
| Streptococcus_sp._I-P16 | 26 | LMAT-ML+H | 7223 | 0.896 |
| Lactococcus_lactis | 25 | LMAT-ML+H | 690 | 0.991 |
| Aggregatibacter_actinomycetemcomitans | 23 | LMAT-ML+H | 1095 | 0.883 |
| Actinomyces_urogenitalis | 22 | LMAT-ML+H | 448 | 0.616 |
| Streptococcus_sp._oral_taxon_056 | 21 | LMAT-ML+H | 122 | 0.656 |
| Streptococcus_sp._HSISS2 | 20 | LMAT-ML+H | 4621 | 0.929 |

\*CP=Clinical Pathoscope, G=GOTTCHA, K=Kraken, M=Metaphlan2

### Detecting Eukaryotes

The LMAT databases (all variants) are the only metagenomic analysis tools that include Eukaryotic sequences in addition to human in the reference database. As a result, we were able to classify reads matching fungi, protozoa, plants, and animals (Table S1). The majority of eukaryotic reads were human, followed by Fungi phylum Dikarya, in which many genera and species were detected. Reads were detected from dog (*Canus, Canus lupus*, or *Canus lupus* domesticus; 3 samples with 49-578 reads each). The fungal genus *Malassezia* had particularly large numbers of reads in many samples, an expected result for a genus naturally found on the skin of many animals. Small numbers of reads for various pathogenic protozoa in the phylum *Apicomplexa* were detected, including *Plasmodium, Acanthamoeba castellanii, Eimeria* (coccidiosis in livestock), *Hammondia hammondi*, and *Toxoplasma gondii*. Five samples contained unusually high numbers of reads (~20,000) of *Hammondia hammondi*, which relies on cats as its definitive host. One stool sample had ~2,700 reads of *Blastocystis hominis*, a gastrointestinal parasite of disputed pathogenicity. Several samples contained 1000-3000 reads of *Entamoeba nuttalli*, known to cause illness in non-human primates. The freshwater ciliated protozoan *Tetrahymena thermophila* was detected in a number of samples, most often with low read count, except a few cases with thousands of reads. One retroauricular crease sample had ~9,500 reads classified as <u>*Trypanosoma cruzi*</u> (Chagas disease or sleeping sickness), which can persist unnoticed in the host for decades, although detection on skin may not mean the host is infected. Small numbers of reads of *Trichomonas vaginalis* were detected in a handful of samples. While the bulk of calls were bacterial, these observations suggest that further study confirming the present of eukaryote reads in the HMP data is warranted. Many draft eukaryotic sequences appear to contain misassembled human and vector/synthetic sequence, and we have invested substantial effort to control and correct for this contamination [14]. We conducted spot checks with BLAST on reads assigned to eukaryotes (including *Malassezia* and *Canus lupus*) to confirm that obvious miss-assignment errors were not present. Nevertheless, non-human eukaryote-classified reads deserve an extra measure of caution, such as demanding higher read count or score thresholds for calling a species as present. LMAT-ML+H is less sensitive than LMAT Grand for these organisms, as manual comparisons of several of these species showed fewer reads and lower scores in the ML, but it is still capable of detecting these organisms using 16-24 GB memory.

## Discussion

Sensitivity was too low for SIANN to be feasible for clinical samples, and false positives were seen in samples with pathogen at high spiked concentrations. MiniKraken processed metagenome reads the fastest, but it was also the least accurate on the real world HMP samples, with poor BLAST support for the species it commonly detected and a large number of missed species that did have strong BLAST support, and classified an average of about a third of the reads. Clinical Pathoscope and GOTTCHA also had poor accuracy, they were the slowest classifiers in our tests, and they failed to classify even larger percentages of reads. MetaPhlan2 was faster than LMAT-ML, although it only classified an average of 5% of reads and missed many species that were clearly present based on BLAST results. LMAT-ML ran on average at about half the speed of MetaPhlan2 and required 24 GB of DRAM to avoid any performance penalty for paging the database index, more than other marker library methods. However, LMAT-ML+H classified over 60% of reads and showed far better accuracy than other marker library methods as verified by BLAST, delivering results nearly as complete as those of LMAT Grand with only a fraction of the memory requirements, at a speed capable of analyzing a gigabase-sized sample in about 3.5 minutes with 24 CPU and 24 GB of DRAM. Additionally, the performance penalty for 16 GB versus 24 Gb is roughly 50% slower with the smaller available DRAM (when using LMAT-ML+H).

The processing rate of LMAT-ML databases are affected by two key factors, the rate at which the index can be accessed, and the classification rate of an individual read, which is impacted by the number of taxonomy identifiers retrieved for each constituent *k*-mer in the read. Fast access to the index is impacted by the size of the database, with larger databases requiring additional paging.

We consider the use of the SATA II SSD as it provides a relatively low cost alternative to DRAM for storage. Current advertised prices for DRAM are about $11 and for SATA SSD $0.75 per gigabyte, and mid tier PCIe flash around $5/GB. We present the use of an SSD with LMAT and limited memory in contrast to previous experiments with PCIe flash with larger LMAT database indexes [19]. Additional experimentation with the larger Grand database and SATA SSD with limited main memory have demonstrated subpar performance. For the smaller ML databases, however, the performance reduction with SATA SSD was minor, since demands on main memory were much lower. Although LMAT runs faster in DRAM only, we estimate that a laptop/desktop with 16 GB RAM and a 24 GB flash drive will perform rapid and accurate metagenome analyses with LMAT-ML+H.

While all methods aim to classify reads at the most specific level possible, that level of specificity must be supported by the data. All of the methods other than LMAT failed to identify genus, family, phylum, or even higher levels of conservation in the HMP reads, and thus reported overly specific calls. Databases like RefSeq have only one representative sequence for many species and some genera, and documentation explicitly states that more than one strain will be included only in exceptional circumstances as determined manually by NCBI staff [20]. This renders suspect any strain calls made by classifiers relying on Refseq, since there are not enough near neighbors to resolve at this level. In addition, MiniKraken, Clinical Pathoscope, and GOTTCHA made errors by misclassifying reads as microbial that were much more likely to be artificial sequences from sample preparation. Metaphlan2 did not misclassify artificial sequences, but it did misclassify human reads as viral. We and others [9, 21] have found substantial contamination of draft genomes with adaptor, vector, and other synthetic sequences and human or other host sequences. LMAT-Grand and LMAT-ML specifically label synthetic and human *k*-mers and then apply a greedy strategy to detect reads with these sequences. This allows the LMAT database to contain large numbers of draft sequences to span novel strain and species diversity without misclassifying human and synthetic reads as microbial due to contaminated draft assemblies.

MiniKraken, Clinical Pathoscope, and GOTTCHA reference sequences consist of NCBI RefSeq complete bacterial, archaeal, viral genomes and the human reference genome, and Metaphlan2 and SIANN use even smaller subsets of these sequences. With LMAT-Grand and LMAT-ML, we extend the reference database to span every microbial genome in the public domain, and more, so that LMAT is the only metagenome classification software that includes 1) eukaryotic sequence in both the Grand and the LMAT-ML databases, enabling the classification of fungi, protozoa, and some multicellular organisms (from organelles labeled as whole genome, e.g. mitochondria and chloroplasts); 2) draft genomes and assembled contigs not contained in NCBI RefSeq; and 3) draft and finished bacteria, virus, archaea, fungi, and protozoa genomes from a number of sequencing centers worldwide with publicly available sequence data in addition to those from NCBI. While sequences available from these sites eventually appear in NCBI databases, they may be publicly available years before release at NCBI, and many strains may never become a part of NCBI RefSeq.

Since LMAT includes all available strains (genomes) that have been sequenced, it should provide more accurate species resolution and reporting of the closest strains in the database than other methods. LMAT reports the number of reads matching multiple strains, and for those reads conserved across multiple strains, then it reports only the species level match, since NCBI taxonomy nodes in general do not exist for clades. Even for isolates with sequenced genomes, there may be more than one best strain match for different subsets of reads due to evolution from the original sequenced isolate during propagation, lateral gene transfer, recombination, and sequencing error. A phylogenetic approach is probably necessary to accurately place new sequence in the broader context of other isolates, possibly using assembly, alignment or SNPs, provided there is sufficient genome coverage. Other methods that fail to include many species and strains in their reference database cannot resolve specific strains. Results presented here show that these methods are incorrect or overly specific even in their species and genus level classifications.

Not only is LMAT the only method that can detect eukaryotic sequences, it reports calls to plasmids versus chromosomes for distinguishing the presence of these mobile genetic elements. Metaphlan2, GOTTCHA and MiniKraken do not detect plasmids, and make only taxonomic calls. Clinical Pathoscope does identify reads by database entry, so it is possible to distinguish plasmid from chromosomal matches. LMAT distinguishes plasmid calls, and creates a file exclusively listing the plasmids detected, and also includes those plasmid calls in the overall results summary with all taxonomic calls. For methods other than LMAT, there is no process described in the manuals for extracting reads responsible for a given call, making it difficult to verify those calls, and do additional analyses such as assembly, per species gene annotation, SNP analyses, or distinguish matches to plasmid versus chromosome. Plus, failure to report standardized NCBI taxonomy identifiers for all calls by some of the methods, plus use of nonstandard or outdated species names and GI numbers not in the current NCBI database makes the process especially challenging. We encourage software developers to describe procedures for extracting reads for the taxonomic calls made by the method, to facilitate call verification from the reads responsible for each call.

Alignment based methods (e.g. BLAST and read mapping) scale linearly with the number of bases in the reference database. To scale with an ever growing pool of reference genomes, alignment-based software must reduce to only a subset of the available data by excluding strain variants, draft genomes, and non-microbial kingdoms. As a consequence, these methods fail to classify large numbers of reads, report overly specific classifications for sequences which in fact are more widely shared across taxa, and either misclassify or fail to detect all the species and genera missing from the database. In addition, alignments require a cap on the maximum number of alignments to return to retain reasonable run times. We have observed that for highly conserved sequences (like 16S rRNA or housekeeping genes) where there may be thousands of additional unreported matches over that maximum, the sort order of reporting matches can result in biases and overly specific calls for taxa that may not actually be present. In contrast, the *k*-mer based approach has the advantage of retaining and condensing conserved subsequences so that adding related reference genomes increases the database size only for novel *k*-mers and the small increment of adding that genome tag to existing *k*-mers already stored. Thus, the database size grows as a function of sequence diversity, not as a strictly linear increase with the number of bases in the reference database. We have plans to add more eukaryotic genomes to the LMAT database (e.g. mosquitos, nematodes, ticks, plants), to classify more reads from environmental samples, which should be tremendously helpful in fields such as bioenergy, microbial ecology, industrial metagenomics, and environmental biosurveillance. [4] Many eukaryotes have extremely large and repetitive genomes [22], so a *k*-mer that scales with diversity rather that the genome size including more eukaryotes should further reduce the number of unclassified reads, and improve our ability to separate reads for truly novel, unknown microbes for further analysis.

Comparing results from actual HMP samples across 6 metagenome analysis software packages, we found that the LMAT Marker Library “LMAT-ML+H” classified microbial contents most accurately and comprehensively due to its reliance on a reference database 1-2 orders of magnitude larger than that of other software and representing 2-4 times more species. Its speed is competitive with other tools, and although memory demands are higher, they are still well within the price range of a standard desktop machine with 24GB of memory or 16GB memory with a low cost SSD drive.

## Footnotes

1 Sanger Center, J. Craig Venter Institute, Baylor College of Medicine, Washington University in St. Louis, Beijing Genome Institute, Integrated Microbial Genomes, European Molecular Biology Laboratory, etc.

2 ftp://ftp.ncbi.nlm.nih.gov/pub/taxonomy/gi_taxid_nucl.dmp.gz, ftp://ftp.ncbi.nih.gov/pub/taxonomy/taxdump.tar.gz, ftp://ftp.ncbi.nlm.nih.gov/gene/DATA/gene2accession.gz

## Acknowledgements

We thank Marisa Torres and Clinton Torres for building the infrastructure to download and update the reference sequence database used by LMAT. This work was performed under the auspices of the US Department of Energy by Lawrence Livermore National Laboratory under Contract DE-AC52-07NA27344. Laboratory Directed Research and Development (33-ER-2012 and 08-ER-2011);

*Conflict of Interest:* none declared.

